# Paired Immunoglobulin-like Type 2 Receptor Alpha G78R variant alters ligand binding and confers protection to Alzheimer’s disease

**DOI:** 10.1101/325936

**Authors:** Nisha Rathore, Sree Ranjani Ramani, Homer Pantua, Jian Payandeh, Tushar Bhangale, Arthur Wuster, Manav Kapoor, Yonglian Sun, Sharookh Kapadia, Lino Gonzalez, Ali A. Zarrin, Alison Goate, David Hansen, Timothy W. Behrens, Robert R. Graham

**Affiliations:** Department of OMNI Human Genetics, Genentech Inc., South San Francisco CA 94080; Department of Microchemistry, Proteomics & Lipidomics, Genentech Inc., South San Francisco CA 94080; Department of Immunology and Infectious Diseases, Genentech Inc., South San Francisco CA 94080; Department of Structural Biology, Genentech Inc., South San Francisco CA 94080; Ronald M. Loeb Center for Alzheimer’s disease, B1065, Icahn School of Medicine at Mount Sinai, 1425 Madison Ave, New York, NY 10029; Department of Immunology, Genentech Inc., South San Francisco CA 94080; Department of Proteomics & Biological Resources, Genentech Inc., South San Francisco CA 94080. Current affiliation: Therapeutics, 23 and Me, South San Francisco, CA 94080; Department of Neuroscience, Genentech Inc., South San Francisco CA 94080; Department of Bioinformatics and Computational Biology, Genentech Inc., South San Francisco CA 94080

## Abstract

Paired Immunoglobulin-like Type 2 Receptor Alpha (PILRA) is a cell surface inhibitory receptor that recognizes specific *O*-glycosylated proteins and is expressed on various innate immune cell types including microglia. We show here that a common missense variant (G78R, rs1859788) of PILRA is the likely causal allele for the confirmed Alzheimer’s disease risk locus at 7q21 (rs1476679). The G78R variant alters the interaction of residues essential for sialic acid engagement, resulting in >50% reduced binding for several PILRA ligands including a novel ligand, complement component 4A, and herpes simplex virus 1 (HSV-1) glycoprotein B. PILRA is an entry receptor for HSV-1 via glycoprotein B, and macrophages derived from R78 homozygous donors showed significantly decreased levels of HSV-1 infection at several multiplicities of infection compared to homozygous G78 macrophages. We propose that PILRA G78R protects individuals from Alzheimer’s disease risk via reduced inhibitory signaling in microglia and reduced microglial infection during HSV-1 recurrence.

## Author summary

Alzheimer’s disease (AD) is a devastating neurodegenerative disorder resulting from a complex interaction of environmental and genetic risk factors. Despite considerable progress in defining the genetic component of AD risk, understanding the biology of common variant associations is a challenge. We find that PILRA G78R (rs1859788) is the likely AD risk variant from the 7q21 locus (rs1476679) and PILRA G78R reduces PILRA endogenous and exogenous ligand binding. Our study highlights a new immune signaling axis in AD and suggests a role for exogenous ligands (HSV-1). Further, we have identified that reduced function of a negative regulator of microglia and neutrophils is protective from AD risk, providing a new candidate therapeutic target.

## Introduction

Alzheimer’s disease (AD) results from a complex interaction of environmental and genetic risk factors [1]. Proposed environmental risk factors include a history of head trauma [2–4] and infection [5–7]. In recent years, large-scale genome-wide association studies (GWAS) and family-based studies have made considerable progress in defining the genetic component of AD risk, and >30 AD risk loci have been identified [8–20].

A key role for microglial/monocyte biology in modulating risk of AD has emerged from analysis of the loci associated with AD risk. Rare variants of TREM2, a microglial activating receptor that signals through DAP12, greatly increase AD risk [11,14]. Beyond TREM2, a number of the putative causal genes mapping to AD risk loci encode microglial/monocyte receptors (complement receptor 1, CD33), myeloid lineage transcription factors (SPI1), and other proteins highly expressed in microglia (including ABI3, PLGC2, INPP5D, and PICALM).

## Results

### PILRA G78R is associated with protection from AD

The index variant for the Alzheimer’s disease risk locus at 7q21 is rs1476679 (meta P value = 5.6 × 10^−10^, odds ratio = 0.91)[15]. In addition to reduced disease risk, the C allele of rs1476679 has been associated with age of onset [21] and lower odds of pathologic AD (plaques and tangles) in the ROSMAP study [22]. In the 1000 Genomes project CEU population (phase 3 data), there were 6 variants with an r^2^>0.9 with rs1476679 (table S1). None of the 6 variants were predicted to alter regulatory motifs that might influence gene expression (Regulome DBscore ≤ 4), but one variant (rs1859788) encoded a missense allele (G78R, ggg to agg transition) in Paired Immunoglobulin-like Type 2 Receptor Alpha (PILRA) protein. Using a cohort of 1,357 samples of European ancestry whole genome-sequenced to 30X average read-depth (Illumina), we confirmed the strong linkage between rs1476679 (in *ZCWPW1* intron) and rs1859788 (G78R PILRA variant) (table S1).

We hypothesized that PILRA G78R was the functional variant that accounts for the observed protection from AD risk. As expected from the strong linkage disequilibrium (LD) between PILRA G78R and rs1476679 (Fig. 1A), conditional analysis demonstrated that the 2 variants were indistinguishable for AD risk in individuals of European ancestry. In a cohort of 8060 European ancestry samples (a subset of samples described in *19*), individuals homozygous for R78 (OR=0.72) and heterozygous (OR=0.89) for R78 were protected from AD risk relative to G78 homozygotes. We note that the allele frequency of PILRA G78R varies considerably in world populations. Indeed *PILRA R78* is the minor allele in populations of African (10%) and European descent (38%) but is the major allele (65%) in East Asian populations [23]. In several non-European populations, rs1476679 (Index risk variant) and rs1859788 (PILRA G78R) are not in tight LD and it may be possible to distinguish the impact on disease risk for these variants. In a dataset of 894 AD cases and 951 controls of Japanese ancestry [24], PILRA G78R conferred an effect size of −0.07 (rs1859788, A allele, P=0.35), while rs1476679 had an effect size of −0.04 (T allele, P=0.65, table S2), suggesting that PILRA G78R associates with AD protection more than rs1476679, although a larger sample is needed for statistical significance. In APOE4 carriers, there was a significant interaction with PILRA G78R (rs1859788, P=0.04), while the interaction of APOE4 and rs1476679 was not significant (P=0.59), further supporting the hypothesis that PILRA G78R may be a casual variant in the 7q21 risk locus (table S2).

The index variant in the 7q21 locus (rs1476679) has been associated with expression levels of multiple genes in the region, including *PILRB* [25,26]. However, a haplotype tagged by rs6955367 is more strongly associated with expression in whole blood of multiple genes in the region (*PILRB, STAG3L5, PMS2P1, MEPCE*), and is only modestly linked to rs1476679 (r^2^=0.085, D’=0.982) in Europeans [27]. Since the PILRB eQTL P value for rs1476679 is not significant (P=0.31) after conditioning rs6955367 (table S3) in whole blood, we conclude that rs1476679 and rs1859788 are not significant causal eQTLs in the 7q21 region that the observed relationship of these SNPs with *PILRB* expression is due to the weakly linked variant rs6955367 (fig. S1). Of interest, the G allele of rs6955367 (increased expression of PILRB) is linked to rs7803454 (r^2^ = 0.83), a variant associated with increased risk of age-related macular degeneration and suggests the presence of independent effects in the PILRA/PILRB region [28].

### PILRA G78R reduces ligand binding

Paired activating/inhibitory receptors are common in the immune system, with the activating receptor typically having weaker affinity than the inhibitory receptor toward the ligands. PILRA and PILRB are type I transmembrane proteins with highly similar extracellular domains that bind certain O-glycosylated proteins [29–32], but they differ in their intracellular signaling domains [33–35]. PILRA contains an immunoreceptor tyrosine-based inhibitory motif (ITIM), while PILRB signals through interaction with DAP12, which contains an immunoreceptor tyrosine-based activation motif (ITAM). Analysis of PILRA knockout mice suggests that PILRA is a negative regulator of inflammation in myeloid cells [36–38], with knockout macrophages showing increased production of cytokines (IL6, IL-1b, KC, MCP-1) in addition to increased infiltration of monocytes and neutrophil via altered integrin signaling.

PILRA is known to bind both endogenous (including COLEC12, NPDC1, CLEC4G, and PIANP) and exogenous ligands (HSV-1 glycoprotein B (gB)) [31,32,37,39]. Because the G78R (R78 (AD protective)) variant resides close to the sialic acid-binding pocket of PILRA, we tested whether the glycine (uncharged, short amino acid) to arginine (basic, long side chain amino acid) substitution might interfere with PILRA ligand-binding activity. All non-human *PILRA* sequences, as well as all *PILRB* sequences, encode glycine at this position. We also generated amino acid point variants in and around the sialic acid-binding pocket of PILRA. A residue conserved among PILR proteins and related SIGLEC receptors, R126 in PILRA, is well known to be essential for sialic acid interaction [30,32,39] and so was not further studied here. Based on their location in the crystal structure, evolutionary conservation [32], and involvement in binding HSV-1 gB [39], amino acids R72 and F76 were predicted to be important for ligand binding and were substituted to alanine as positive controls for loss-of-function [32]. In addition, S80, a residue outside of the sialic acid-binding pocket was substituted to glycine. The R72A, F76A, and S80G mutations have not been detected in human populations (dbSNP v147).

To study receptor-ligand binding, 293T cells were transfected with G78 (AD risk) PILRA or variants, and then incubated with purified NPDC1-mIgG2a protein (Fig. 1B), followed by flow cytometry to detect PILRA and the NPDC1 fusion protein. Among known PILRA ligands, NPDC1 is expressed in the central nervous system and binds with high affinity to PILRA [32]. Expression of the PILRA variants on the transfected 293T cells was comparable to or greater than G78 (AD risk) PILRA (fig. S2). G78 (AD risk) PILRA binding to NPDC1 was considered 100%. Both R72A and F76A mutations severely impaired NPDC1 binding (~20% of G78, p-value < 0.0001). The R78 (AD protective) variant also showed significantly reduced ligand binding (~35% of G78, p < 0.0005), while the G80 mutant was the least affected (~60% of G78, p < 0.0001) (Fig. 1c and fig. S3a,b).

To further test the hypothesis that the AD protective PILRA R78 variant impacts ligand binding, NPDC1 or alternative PILRA ligands HSV-1 gB and PIANP were expressed on the cell surface of 293T cells, and the binding of purified PILRA protein variants was measured by flow cytometry. PILRA R78 showed reduced binding to the various ligands in these assays as compared to G78 (Fig. 1, D to G and fig. S4, A to G). These data confirmed that the R78 variant impairs ligand-binding activity of PILRA.

**Figure 1.**
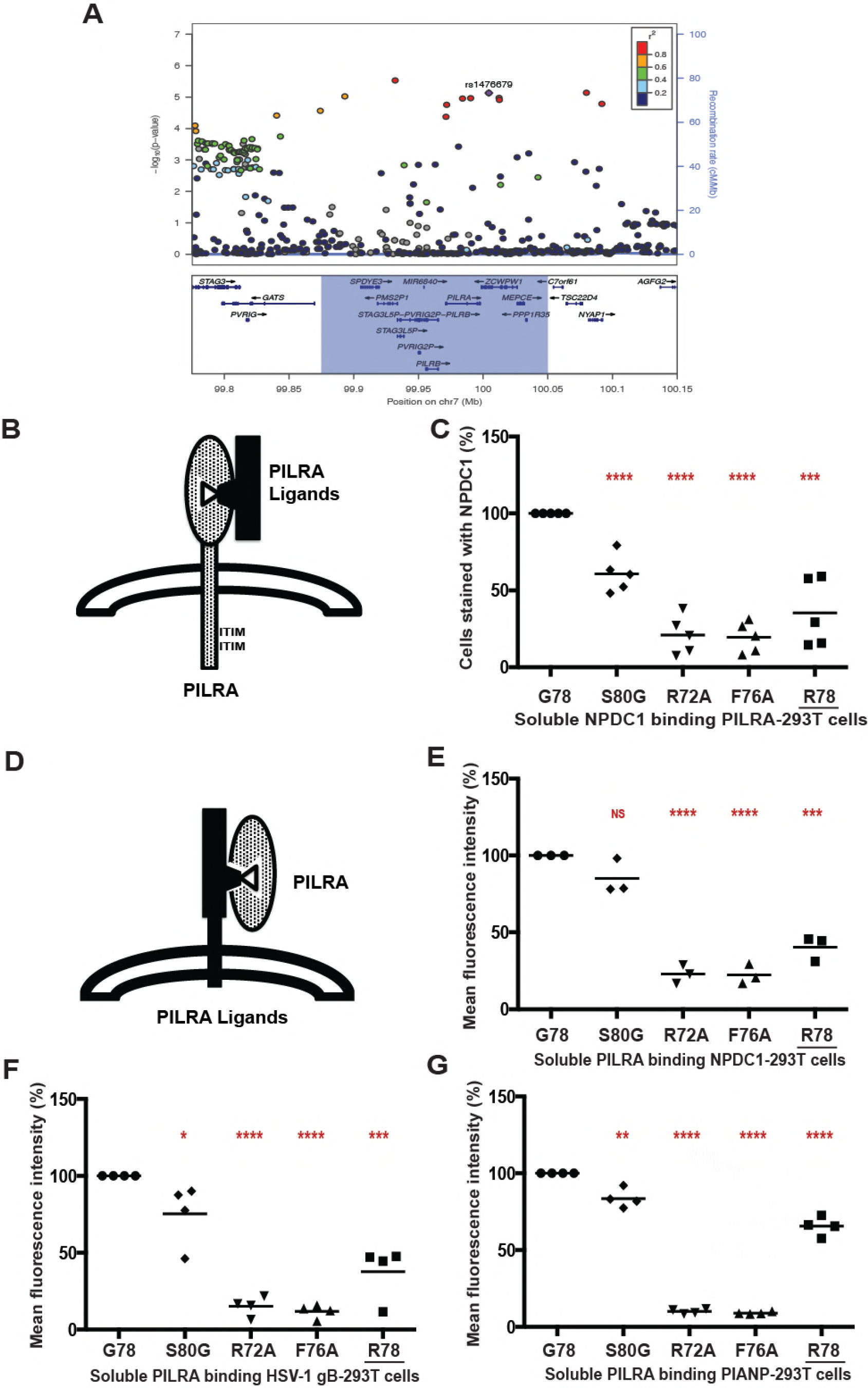
PILRA G78R reduces ligand binding. A) Association of variants in the 7q21 locus with AD risk in the IGAP phase 1 dataset [15]. B,C) 293T cells were transfected with G78 (AD risk) PILRA and several point mutants of PILRA. Binding of NPDC1-mFC to PILRA variant-transfected cells was measured by flow cytometry. The percent of cells expressing PILRA and positive for NPDC1 is indicated in each *panel* considering G78 (AD risk) PILRA binding as 100% for each experiment D,E,F,G) In the inverse experiment, 293T cells were transfected with different known ligands of PILRA (NPDC1, HSV-1 gB and myc-PIANP). Binding of different PILRA variants to ligand-transfected cells was analyzed by flow cytometry. Results are the percentage of MFIof PILRA-mFC binding on ligand-transfected cells considering G78 (AD risk) PILRA binding as 100% for each experiment. Statistical analysis is two-tailed unpaired t-test (p values <0.05= *, <0.005=**, <0.0005=***, <0.0001=****) performed on 3-5 independent experiments.

### Identification of C4A as PILRA ligand

A peptide motif for PILRA interaction has been established (Fig. 2A) that includes an O-glycosylated threonine, an invariant proline at the +1 position, and additional prolines at the −1 or −2 and +3 or +4 positions [32,39]. Of note, PILRA is capable of binding murine CD99 and human NPCD1 (both contain the consensus motif), but not human CD99 or murine NPCD1 (both lack the consensus motif), suggesting divergence between human and mouse in the range of endogenous ligands bound by PILRA [32].

We sought to identify novel endogenous PILRA ligands by searching for human proteins with either the PTPXP, PTPXXP, PXTPXP or PXTPXXP motif. A total of 1540 human proteins carry at least 1 of these putative PILRA-binding motifs (table S4). Narrowing the search, we considered proteins with the motif that have previously been shown to be O-glycosylated in human cerebral spinal fluid [40], and measured the binding of these proteins to PILRA variants. By flow cytometry, complement component 4A (C4A) bound to G78 (AD risk) PILRA in a manner comparable to NPDC1, while APLP1 and SORCS1 showed relatively little interaction with PILRA (Fig. 2B and fig. S5, A and B). We further demonstrated that the PILRA R78 (AD protective) variant has reduced binding for C4A (Fig. 2C and fig. S5C). We did not test C4B, but its putative PILRA-binding motif is identical to that of C4A.

**Figure 2.**
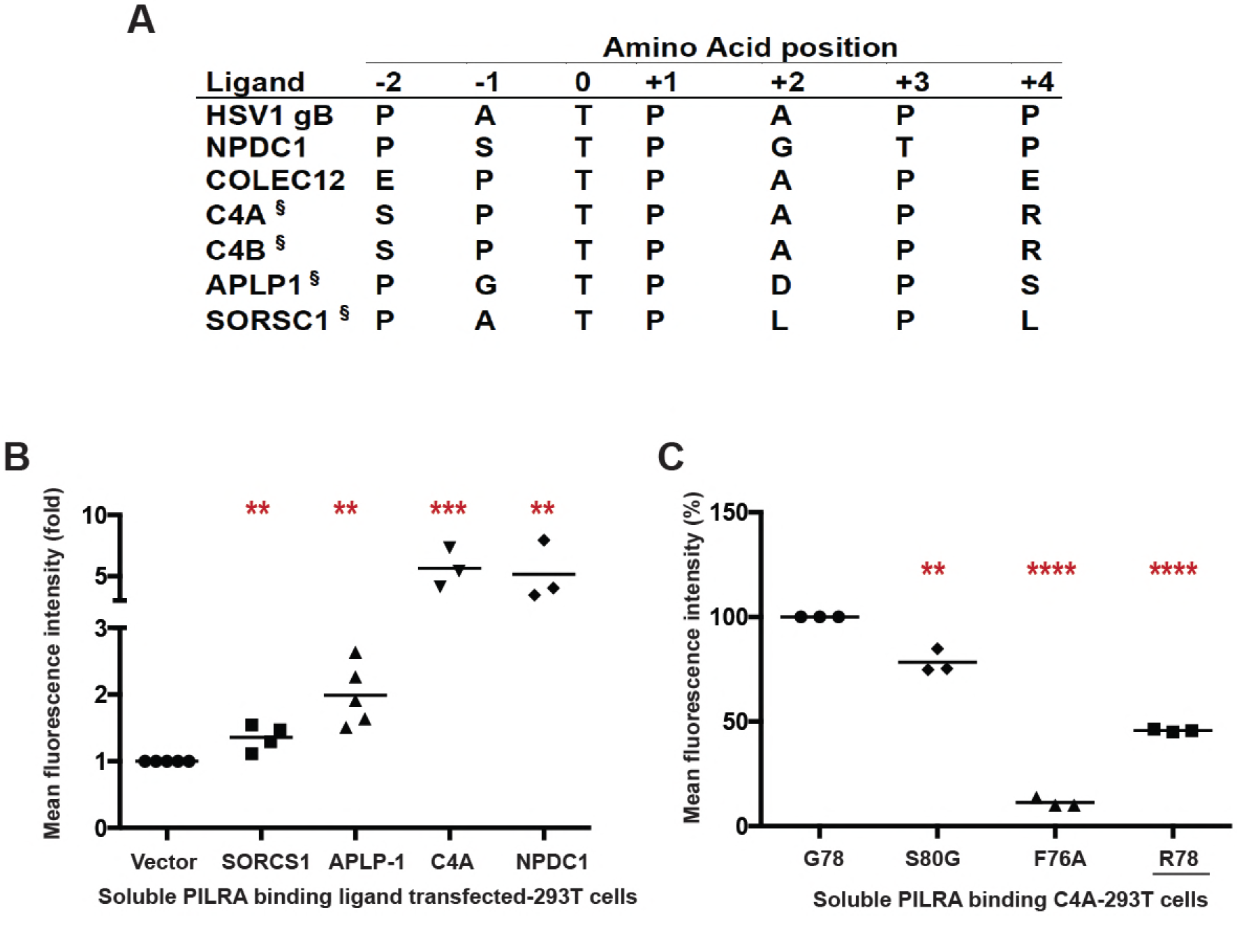
C4A is a novel ligand for PILRA. A) Comparison of the peptide sequence around the O-glycosylated Thr (position 0) of known and putative (§) PILRA ligands. B)293T cells were transfected with putative ligands of PILRA (SORCS1 ECD, APLP1 ECD. or full length C4A) fused with C-terminal glycoprotein D (gD) tag and GPI anchor, or full length NPDC1 as positive control. Binding of G78 (AD risk) PILRA to ligand-transfected cells was analyzed by flow cytometry. Results are fold-increase in binding to each putative ligand compared to vector control for each experiment. C) 293T cells were transfected with full length C4A fused with C-terminal gD tag and GPI anchor. Binding of different PILRA variants to C4A-transfected cells was analyzed by flow cytometry. Results are the percentage of MFIof PILRA-mFc binding on ligand-transfected cells considering G78 (AD risk) PILRA binding as 100% for each experiment. Statistical analysis is two-tailed unpaired t-test (p values <0.05= *, <0.005=**, <0.0005=***, <0.0001=****) performed on 3-5 independent experiments.

### G78R stabilizes the ligand-free state of PILRA

To understand the conformational changes that might occur in the PILRA sialic acid-binding pocket during receptor-ligand interactions in the presence of G78 (AD risk) or R78 (AD-protective) variants, we evaluated available experimental crystal structures (Fig. 3, A to C) [39,41]. Structures of G78 (AD risk) PILRA reveal a monomeric extracellular domain with a single V-set Ig-like β-sandwich fold that binds O-glycan ligands (Fig. 3, B and C) [39]. By analogy to a molecular clamp, the sialic acid-binding site in PILRA undergoes a large structural rearrangement from an “open” to a “closed” conformation upon binding its peptide and sugar ligands simultaneously (Fig. 3, A to C). The essential R126 side-chain engages the carboxyl group of sialic acid (SA) directly in a strong salt bridge (Fig. 3C). The CC’ loop which contains F76 and G78 undergoes a large conformational change where F76 translates ~15 Å to participate in key interactions with the peptide of the ligand and abut the Q140 side-chain of PILRA (Fig. 3, B and C). In this ligand-bound “closed” conformation of PILRA, Q140 helps to position R126 precisely for its interaction with SA (Fig. 3C).

Notably, in the structure of R78 (AD protective) PILRA crystallized in the absence of any ligand [41], the long side-chain of R78 is observed to hydrogen bond with Q140 directly (Fig. 3A). This unique R78-Q140 interaction has three important consequences: 1) it sterically hinders F76 from obtaining a ligand-bound “closed” conformation, 2) it affects the ability of R126 to interact with the carboxyl group of SA by altering the R126-Q140 interactions observed in G78 (AD risk) PILRA and, 3) it likely alters CC’ loop dynamics, (Fig. 3, B to C). Overall, the structure of the R78 (AD protective) variant shows that this single side-chain alteration appears to stabilize the “open” apo form of PILRA and likely alters the conformational sampling of the molecular clamp required to obtain its “closed” form to engage its ligands.

**Figure 3.**
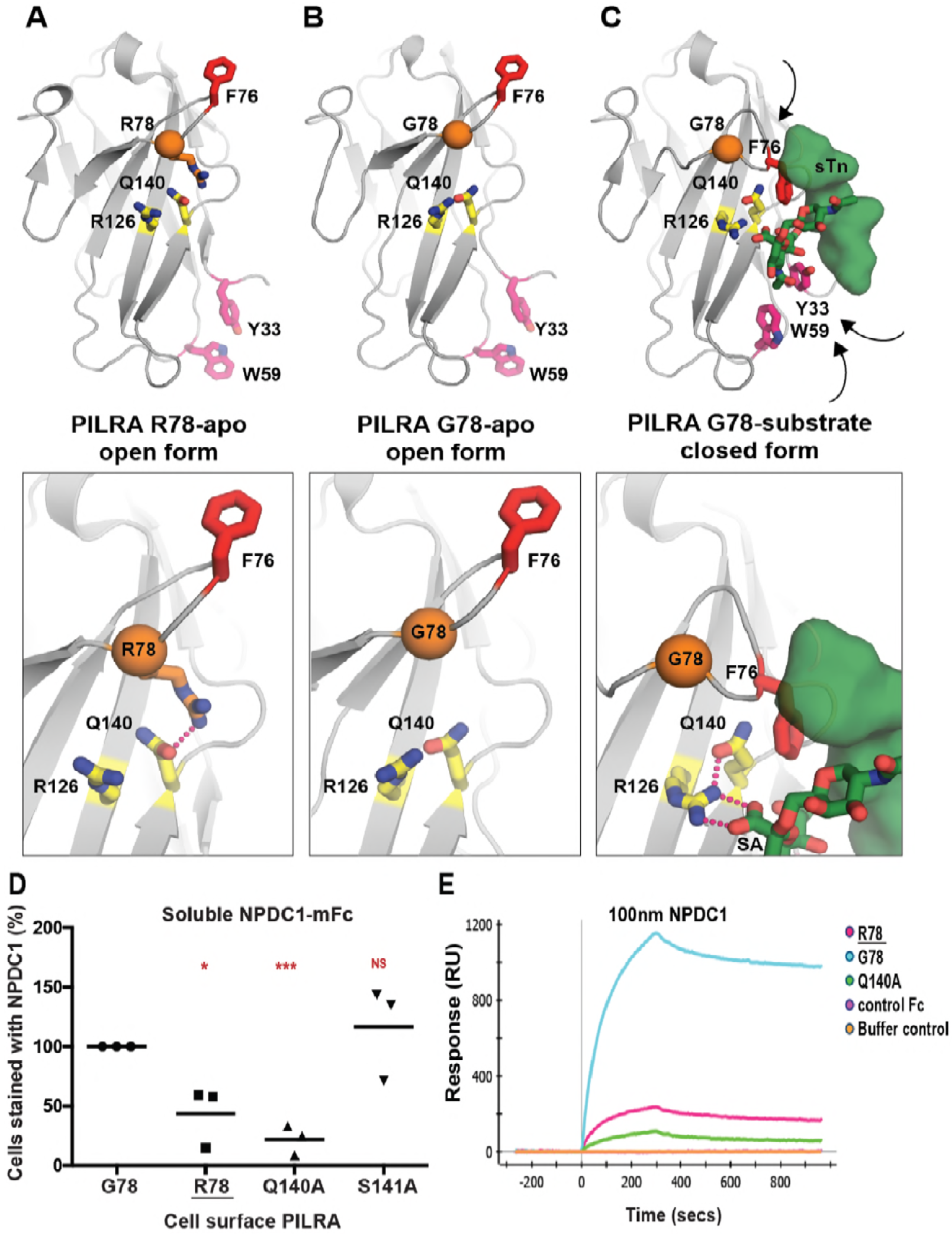
Structural determinants of PILRA in apo and ligand-bound conformations. A) The unliganded crystal structure of R78 (AD protective) PILRA (PDB 4NFB) displays an “open” conformation with an unformed sialic acid (SA) binding site. B) The apo crystal structure of G78 (AD risk) PILRA (PDB 3WUZ) reveals that the Q140-R126 interaction network (productive for SA-coordination) is pre-formed and the “downward” movement of F76 is not impeded in the R78 side chain mediated interactions. C) The sialylated *O*-linked sugar T antigen (sTn)-bound PILRA structure (PDB 3WV0) reveals the ligand-induced conformational changes across the receptor that lead to engagement of the SA-motif by direct coordination to R126 and the critical involvement of F76 in peptide (sTn ligand) recognition. Aromatic residues including Y33 (fuchsia) and W59 (fuchsia) also undergo significant ligand-induced conformational changes. D) 293T cells were transfected with G78 (AD risk) PILRA and several point mutants of PILRA. Binding of NPDC1-mFC to PILRA variant-transfected cells was measured by flow cytometry. The percent of cells expressing PILRA and positive for NPDC1 is indicated in each *panel* considering G78 PILRA binding as 100% for each experiment. Statistical analysis is two-tailed unpaired t-test. E) Binding of NPDC1.mFC to PILRA variants by surface plasmon resonance (SPR).

We therefore propose that in G78 PILRA (AD-risk associated), the engagement of SA by R126 and peptide by F76 is facilitated by G78 (Fig. 3C). However, in the AD-protective PILRA variant R78, the R78 side-chain competes with the central R126-Q140 interaction and alters the positioning of F76 (Fig. 3A), which leads to an overall decrease in PILRA ligand binding. This structure-based hypothesis is consistent with the reduced functional cellular binding observed for the R78 variant (Fig. 1).

To further test this model, we generated two additional alanine mutants of PILRA at amino acids predicted to be essential (Q140) or non-essential (S141) for conformational changes associated with ligand interaction. 293T cells were transfected with G78 (AD risk), R78 (AD protective), Q140A and S141A variants of PILRA, and receptor-ligand interaction was measured after incubating cells with soluble NPDC1-mIgG2a. PILRA expression was comparable among variants, matching or exceeding G78 (AD risk) expression (fig. S2). R78 (44% of G78, p=0.02) and Q140A (22% of G78, p=0.0004) variants showed significantly decreased binding to NPDC1, while S141A (117% of G78, p=0.5) had no significant effect (Fig. 3D and fig. S6, A and B). These data are consistent with the experimental structural models that show the interaction of Q140 with R126 is important for productive sialic acid binding (Fig. 3, A to C). Consistently, the Q140A mutation has a strong effect because the Q140-R126 interaction network is completely abolished. By contrast, the AD-protective R78 variant likely has an intermediate effect since it only modulates the Q140 interaction with R126, which is expected to only alter the frequency or strength of relevant PILRA-ligand interactions.

### PILRA G78R reduces the on-rate of ligand binding

We next investigated the interaction of PILRA variant and ligands *in vitro* using surface plasmon resonance (SPR). Human PILRA-Fc variants (G78, R78, or Q140A) were immobilized on a ProteOn GLC sensor chip and binding of NPDC1-mFc or a control mFc-tagged protein was measured. Qualitatively, NPDC1-Fc bound to the R78 (AD-protective) and Q140A (essential for R126 conformation) variants to a much lesser extent than to G78 (AD risk) PILRA, while control Fc-tagged protein showed no binding (Fig. 3E).

To further probe the mechanistic basis of R78 (AD protective) function and phenotype, a more complete SPR characterization of NPDC1-His binding to PILRA variants was performed (fig. S6C). The affinity of NPDC1 toward R78 (AD-protective) PILRA (76.5 nM) was 4.5-fold weaker than the affinity toward G78 PILRA (16.8 nM). The on-rate constant k_on_ for NPDC1-His binding to R78 (AD protective) (6.8×10^+3^ M^−1^s^−1^) was ~3-fold lower than binding to G78 (AD risk) PILRA (2.2×10^+4^ M^−1^s^−1^), while the *k*_off_ rate constants were comparable (fig. S6C). These results are consistent with the idea that, once engaged, the affinity and disassociation rate of R78-ligand complexes are similar to G78 PILRA, but the frequency with which PILRA can productively engage with ligands is reduced in the R78 (AD protective) variant by R78 side chain interactions favoring the apo-state (Fig. 3). Taken together, these data support a structural model in which R78 impairs PILRA-ligand interactions by altering the accessibility of a productive sialic acid-binding conformation in PILRA.

### PILRA G78R reduces entry of HSV-1 into hMDMs

Given that PILRA is a known entry receptor for HSV-1 [42] and the R78 (AD protective) variant showed reduced binding to HSV-1 gB (Fig. 1F), we next determined whether there were differences in HSV-1 infectivity based on PILRA genotype. We isolated and differentiated human monocyte-derived macrophages (hMDMs) from five pairs of healthy volunteers homozygous for either the G78 (AD risk) or R78 (AD protective) PILRA variants (matched for age, gender and ethnicity). hMDMs were infected with HSV-1 at different multiplicities of infection (MOI) (0.01, 0.1, 1 and 10), and infectivity was measured morphologically by light microscopy, by using an LDH cytotoxicity assay, by measuring intracellular viral DNA and in a viral plaque assay.

No notable cytopathic effects were observed in the first 6 h of infection, however at 18 hours post infection, extensive cytopathy was detected in G78/G78 PILRA-expressing hMDMs, including loss of cell shape, increased cell volume, birefringence, and formation of both cell aggregates and multinucleated giant cells (syncytia) (Fig. 4A and fig. S7). Cytopathic changes were less pronounced in R78/R78 (Alzheimer’s protective) homozygous hMDMs (Fig. 4A and fig. S7).

hMDMs from R78/R78 PILRA donors showed significantly less HSV-1-induced cytotoxicity at 18 hrs post infection in the LDH assay at 0.01, 0.1, or 1 MOI (Fig. 4B and table S5). The difference was no longer significant at 10 MOI or if the infection was allowed to proceed for 36 hrs, except at the lowest MOI of 0.01 (Fig. 4B, fig. S8A, and tables S5 and S6).

hMDMs from R78/R78 donors showed 5-10 fold decreased amounts of HSV-1 DNA at 6 hrs at all MOIs (0.01, 0.1, 1 and 10), and at 18 hrs at lower MOIs (0.01 and 0.1), compared to those from G78/G78 donors (Fig. 4C and fig. S8, C and D). No significant differences in HSV-1 DNA were observed between the two genotypes at 18 hrs at higher doses (1 and 10 MOI) (Fig. 4C and fig. S8D), or at 36 hrs for any dose of virus (fig. S8, B and E).

Finally, we measured the amount of infectious HSV-1 virus by harvesting supernatants from HSV-1-infected hMDMs and measuring viral titer by plaque assays on Vero cells. Viral plaque forming units (PFUs) were significantly lower after 6 and 18 hrs of infection for all MOIs tested, and at 36 hrs for lower MOIs (Fig. 4, D and E, and fig. S9). Taken together, these data indicate that R78/R78 macrophages were less susceptible to HSV-1 infection than G78/G78 macrophages.

**Figure 4.**
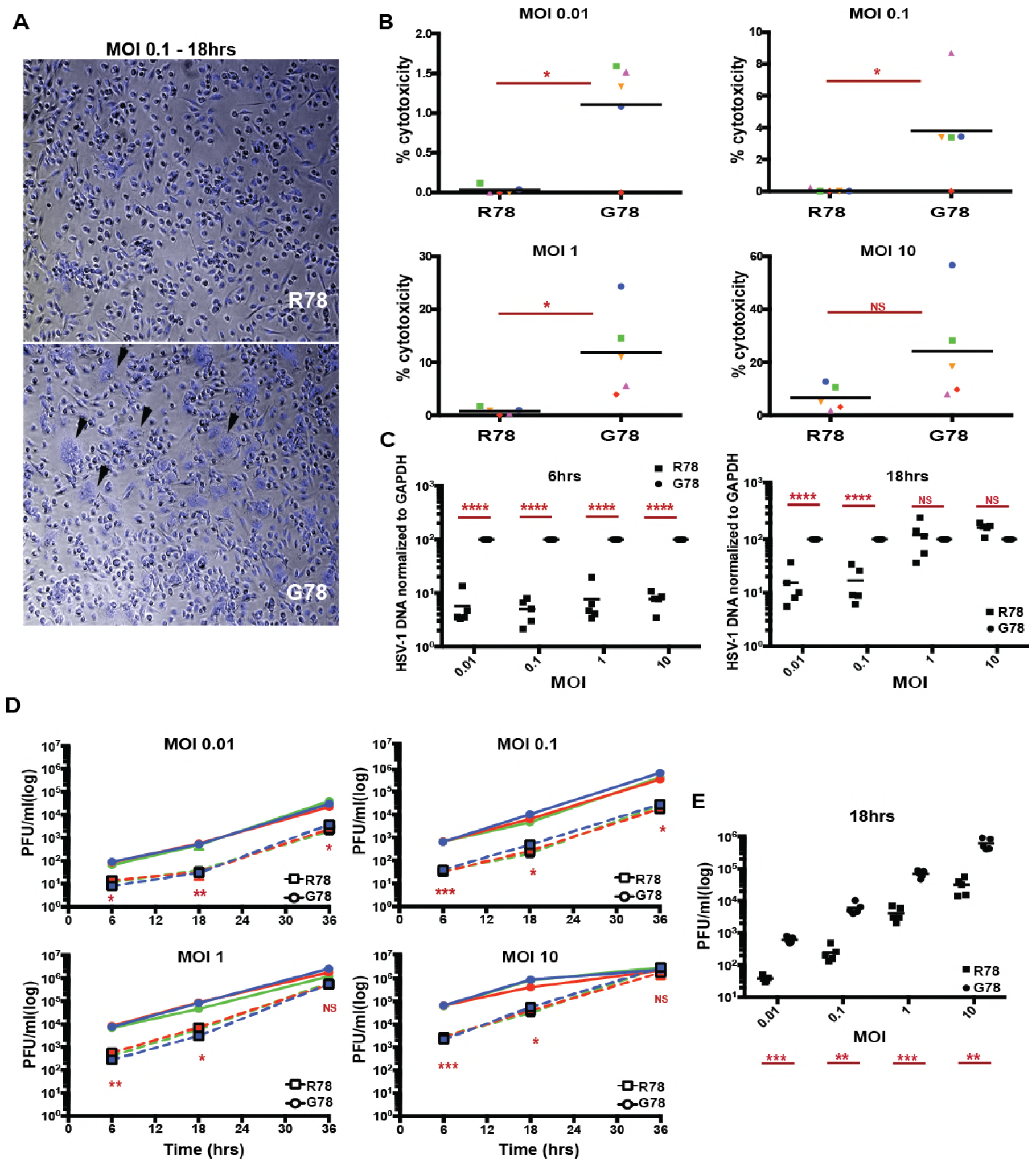
PILRA G78R reduces entry of HSV-1 into hMDMs. A) Representative images of hMDM, infected with HSV-1 for 18 hrs at MOI 0.1, were fixed and stained with DAPI. B) LDH cytotoxicity assay was performed on supernatants harvested from HSV-1-infected hMDMs after 18 hrs. Results are % cytotoxicity - amount of LDH in supernatant after infection compared to LDH released from cells completely lysed by lysis buffer, with completely lysed cells (maximum LDH release) considered as 100% for each donor. Each shape represents one donor pair. C) HSV-1 DNA was quantitated on DNA extracted from HSV-1-infected hMDMs after 6 and 18 hrs by qPCR. Results are % HSV-1 DNA normalized to GAPDH considering G78 donor as 100% for each donor pair. D,E) Viral titers in the supernatant of HSV-1-infected hMDMs were determined by plaque assay on Vero cells. Results are number of plaque forming units (pfu) per ml of supernatant collected from HSV-1-infected hMDMs. (D) 6, 18 and 36 hrs of infection (G78, solid lines; R78, dashed lines)(experiment 1) (E) 18 hrs of infection (data from two individual experiments). Statistical analysis is two-tailed paired (B,D,E) or unpaired (C) t-test performed on 3-5 genotyped individual donor pairs (p values <0.05=*, <0.005=**, <0.0005=***, <0.0001=****)

## Discussion

We show here that PILRA G78R is a likely causal variant conferring protection from AD risk at the 7q21 locus. G78R alters the access to SA-binding pocket in PILRA, where R78 PILRA shows reduced binding to several of its endogenous cellular ligands and with HSV-1 gB. Reduced interaction with one or more of PILRA’s endogenous ligands (including PIANP and NPDC1) could impact microglial migration or activation [36–38]. In fact, microglia up-regulate the expression of the *PIANP* gene in the PS2APP, 5xFAD, and APP/PS1 mouse models of AD [43–45], The identification of C4 as a novel PILRA-interacting protein is also intriguing, given the increased expression of C4 in mouse AD models [43], the increase in amyloid deposition observed when complement activation is inhibited [46], and the genetic association of complement receptor 1 with AD [47]. Finally, we note that both TREM2 and PILRB function as activating receptors and signal through DAP12 [33,35,48]. A reduction of PILRA inhibitory signals in R78 carriers could allow more microglial activation via PILRB/DAP12 signaling and reinforce the cellular mechanisms by which TREM2 is believed to protect from AD incidence [49]. The relevant ligands for PILRA/PILRB *in vivo* and the mechanism by which reducing PILRA-ligand interaction confers protection from Alzheimer’s disease remain to be elucidated.

A role for infection in accelerating AD has been proposed, but remains controversial [50]. HSV-1 is a neurotropic virus that infects a large fraction of the adult population and has frequent reactivation events. HSV-1 has been implicated in AD pathogenesis by several lines of evidence, including the presence of HSV-1 viral DNA in human brain tissue [51,52], increased HSV-1 seropositivity in AD cases [53–56], the correlation of high avidity HSV-1 antibodies with protection from cognitive decline [56], the binding of HSV-1 gB to APOE-containing lipoproteins [57], HSV-1-induced amyloidogenic processing of amyloid precursor protein (APP) [58–60], and preferential targeting of AD-affected regions in HSV-1 acute encephalitis [61]. In addition, HSV-1 gD receptors and gB receptor PILRA increase with age in multiple brain regions, including the hippocampus [62]. Additional AD risk loci have been proposed to play a role in the life cycle of HSV-1 [63], including CR1, which is capable of binding HSV-1 [64]. The reduced infectivity of HSV-1 in R78/R78 macrophages suggests that brain microglia from R78/G78 and R78/R78 individuals are less susceptible to HSV-1 infection and more competent for immune defense during HSV-1 recurrence.

These data provide additional evidence for a key role of microglia in AD pathogenesis and provide a mechanism by which HSV-1 may contribute to AD risk. Inhibiting the interaction of PILRA with its ligands could therefore represent a novel therapeutic mechanism to prevent or slow AD progression.

## Materials and Methods

### PILRA variants and PILRA ligands expression and purification

The coding sequences (CDS) of full length PILRA (AJ400841), human herpesvirus 1 strain KOSc glycoprotein B (HSV-1 gB) (EF157316), and neural proliferation, differentiation and control 1 (NPDC1) (NM_015392.3) were cloned in the pRK neo expression vector. Several PILRA point mutations were generated, including R72A, F76A, G78R, S80G, Q140A and S141A. The PILRA variants were incorporated into a full-length G78 (AD risk) PILRA construct by site-directed mutagenesis as per the manufacturer’s recommendation (Agilent Cat. No. 200523) and sequences were verified. A full length myc-DDK tagged PIANP construct was purchased from Origene (Cat. No. RC207868). Full length complement component 4A (Rodgers blood group) C4A (NM_007293.2), extra cellular domain (ECD) of amyloid beta precursor like protein 1 (APLP1) (NM_005166) (1-580 aa) and ECD of sortilin-related VPS10 domain-containing receptor 1 (SORCS1) (NM_052918) (1-1102 aa) were fused with C-terminal gD tag (US6/gD, partial [Human alphaherpesvirus 1) (<u>AAP32019.1</u>) and GPI anchor in pRK vector. The ECD of all PILRA variants (1-196 aa) and NPDC1 (1-190 aa) were PCR amplified and cloned with C-terminal murine IgG2a Fc tag in a pRK expression vector.

ECDs of PILRA variants (G78 (AD risk), R72A, F76A, G78R, S80G, Q140A and S141A) and NPDC1 fused to the Fc region of murine IgG2a were expressed in a CHO cell expression system, supernatants collected, protein A/G affinity-purified and verified by SDS-PAGE and mass spectroscopy.

### Relative PILRA-ligand binding to PILRA variant transfected cells

293T cells were transfected with lipofectamine LTX reagent (ThermoFisher) with various full-length constructs of PILRA variants (G78 (AD risk), R72A, F76A, G78R, S80G, Q140A and S141A). After 48 hours, the transfected cells were harvested and incubated with soluble mIgG2a-tagged ligand, NPDC1-mFc at 50 ◻g/ml (as described above) for 30 minutes on ice. Cells were then washed and stained with 1 ◻g/ml chimeric anti-PILRA antibody (mouse Fc region is substituted to human IgG1 backbone on anti-PILRA antibodies [32]) on ice for 30 min followed by APC-conjugated mouse anti-human IgG (BD Pharmingen Cat.No. 550931) and FITC anti-mouse IgG2a (BD Pharmingen Cat. No. 553390) secondary antibodies according to manufacturer’s instruction. PILRA-transfected 293T cells were examined by flow cytometery for binding of NPDC1 by measuring the frequency of APC and FITC double-positive cells. Double positive cells were gated on the WT sample and than the gates were overlaid on subsequent samples to maintain the same cell population through out the experiment. For each PILRA variant, the mean percentage of the number of cells binding to NPDC1-mFC relative to the wild type PILRA binding for each experiment was calculated.

### Relative PILRA variant binding to PILRA ligand transfected cells

In the inverse experiment, 293T cells were transfected with lipofectamine LTX reagent [ThermoFisher] with known full-length PILRA ligand (NPDC1, HSV-1gB and PIANP) and predicted ligand constructs (SORCS1, APLP1 and C4A) (described above). After 48 hours, the transfected cells were harvested and incubated with soluble mIgG2a-tagged variants of PILRA (G78 (AD risk), R72A, F76A, G78R, S80G) (described above) 50 ◻g/ml for 30 min on ice. Cells were then washed and stained with FITC anti-mouse IgG2a (BD Pharmingen Cat. No. 553390) secondary antibody according to manufacturer’s instruction. PILRA ligand-transfected 293T cells were examined by flow cytometry for binding to PILRA variants by measuring the frequency of FITC-positive cells. The percentage of mean flouresence intensity (MFI) of PILRA-mFC binding on ligand-transfected cells relative to the wild type PILRA binding for each experiment was calculated.

### PILRA variant ligand binding Surface Plasmon Resonance (SPR)

Binding of human NPDC1.Fc to PILRa-Fc variants was measured by SPR using a ProteOn XPR36 (Bio-Rad). PILRA-Fc WT and variants (G78R and Q140A) were immobilized on a ProteOn GLC sensor chip (Bio-Rad) by EDC/NHS amine coupling (2000-2400 RU’s) and the chip surface was deactivated by ethanolamine after immobilization. NPDC1-Fc diluted in PBST or a control Fc-tagged protein was injected at a concentration of 100 nM over the immobilized PILRA proteins at room temperature[32].

### Isolation and differentiation of monocytes

Healthy human volunteers from the Genentech Genotype and Phenotype program (gGAP) were genotyped for rs1859788 (PILRA G78R) using custom design ABI SNP genotyping assay with the following primers; Forward primer seq: GCGGCCTTGTGCTGTAGAA, Reverse primer seq: GCTCCCGACGTGAGAATATCC, Reporter 1 sequence: VIC-ACTTCCACGGGCAGTC-NFQ, Reporter 2 sequence: FAM-ACTTCCACAGGCAGTC-NFQ. To control for a possible effect of the eQTL for PILRB, all volunteers selected were homozygous AA (lower PILRB expression) for rs6955367 (http://biorxiv.org/content/early/2016/09/09/074450). Genotype for rs6955367 was determined using an InfiniumOmni2.5Exome-8v1-2_A.bpm. Peripheral Blood Mononuclear Cells (PBMC’s) were obtained by Ficol gradient from five pairs of homozygous donors for rs1859788 (one with each genotype AA/GG). The pairs of samples were matched for age [± 5 years], gender and self-reported ethnicity. Monocytes were purified from PBMC’s by negative selection using the EasySep™ Human Monocyte Enrichment Kit without CD16 Depletion (19058), as recommended by the manufacturer. Isolated monocytes were differentiated into macrophages in DMEM + 10%FBS + 1X glutaMax and 100 ng/ml MCSF media for 7-10 days. The gGAP program was reviewed and approved by the Western Regional Institutional Board.

### HSV-1 Infection of Macrophages

Macrophages differentiated from healthy human monocytes were incubated with 10, 1, 0.1 and 0.01 multiplicity of infection (MOI) of HSV-1 virus at 37°C for 1 hour with gentle swirling to allow virus adsorption. Cells were washed after 1 hr of adsorption and infection was continued for 6, 18 and 36 hrs. Supernatant was harvested at 6, 18 and 36 hrs of infection and cell debris were removed by centrifugation at 3000 rpm for 5 min at 4°C. DNA was isolated from infected cells using the QIAamp DNA mini-kit (Qiagen Cat. No. 51304). Additional cells were fixed with 4% paraformaldehyde after infection and stained with DAPI for microscopy.

### Lactate Dehyrogenase (LDH) Cytotoxicity Assay

The CytoTox 96^®^ Non-Radioactive Cytotoxicity Assay (Promega Cat. No. E1780) was performed on supernatant harvested from HSV-1-infected human macrophages as per manufacturer’s recommendations to measure cell toxicity after HSV-1 infection. For each sample, the percent cytotoxicity was calculated as the ratio of LDH released in culture supernatant after infection to completely lysed cells (maximum LDH release).

### Quantitative Polymerase Chain Reaction

HSV-1 DNA was quantitated using a custom design ABI TaqMan gene expression assay, with the following primers: Forward primer seq: 5′-GGCCTGGCTATCCGGAGA-3′, Reverse primer seq: 5′-GCGCAGAGACATCGCGA-3′, HSV-1 probe: 5′-FAM-CAGCACACGACTTGGCGTTCTGTGT-MGB-3′. GAPDH DNA was quantitated using ABI endogenous control (Applied Biosystem Cat. No. 4352934E). Amplification reactions were carried out with 5 ◻L of extracted DNA from infected cells in a final volume of 25 ◻l with TaqMan Universal PCR Master Mix (Applied Biosystems Cat. No. 4304437) as per manufacturer’s recommendations. HSV-1 DNA (Ct values) was normalized to cell GAPDH (Ct values) to account for cell number.

### HSV-1 Plaque Assay

Virus titers from HSV-1-infected cells were determined following a standard plaque assay protocol [65]. In brief, the plaque assay was performed using Vero cells (African Green Monkey Cells) seeded at 1×10^5^ cells per well in 48-well plates. After overnight incubation at 37°C, the monolayer was ~90-100% confluent. Supernatants harvested from HSV-1-infected human macrophages were clarified from cells and debris by centrifugation at 3000 rpm for 5 minutes at 4°C. Virus-containing supernatants were then diluted from 10^−1^ to 10^−8^ in DMEM (1 ml total volume). Growth media was removed from Vero cells and 250 ◻l of supernatant dilution was transferred onto the cells, followed by incubation at 37°C for 2 hrs with gentle swirling every 30 min to allow virus adsorption, after which the virus-containing media was aspirated. The cells were then overlaid with 2% methylcellulose containing 2X DMEM and 5% FBS and incubated at 37°C. 48 hrs post-infection, plaques were enumerated from each dilution. Virus titers were calculated in pfu/ml.

## Acknowledgements

We thank Lindsay Farrer for sharing summary statistics for the PILRA/B region from PLoS One. 2013; 8(4): e58618. The authors would also like to thank the GAP donation program at Genentech and all the donors for facilitating blood samples.

## Supporting information legends

Supplementary Information file contains Supplementary Figures S1-S10 and Supplementary tables S1-S6

## References

1. Holtzman DM, Morris JC, Goate AM. Alzheimer’s Disease: The Challenge of the Second Century. Sci Transl Med. 2011;3: 77sr1–77sr1. doi:10.1126/scitranslmed.3002369

2. O’Meara ES, Kukull WA, Sheppard L, Bowen JD, McCormick WC, Teri L, et al. Head injury and risk of Alzheimer’s disease by apolipoprotein E genotype. Am J Epidemiol. 1997;146: 373–84. Available: http://www.ncbi.nlm.nih.gov/pubmed/9290497

3. Tang MX, Maestre G, Tsai WY, Liu XH, Feng L, Chung WY, et al. Effect of age, ethnicity, and head injury on the association between APOE genotypes and Alzheimer’s disease. Ann N Y Acad Sci. 1996;802: 6–15. Available: http://www.ncbi.nlm.nih.gov/pubmed/8993479

4. Jordan BD, Relkin NR, Ravdin LD, Jacobs AR, Bennett A, Gandy S. Apolipoprotein E epsilon4 associated with chronic traumatic brain injury in boxing. JAMA. 1997;278: 136–40. Available: http://www.ncbi.nlm.nih.gov/pubmed/9214529

5. Kumar DKV, Choi SH, Washicosky KJ, Eimer WA, Tucker S, Ghofrani J, et al. Amyloid-β peptide protects against microbial infection in mouse and worm models of Alzheimer’s disease. Sci Transl Med. 2016;8: 340ra72. doi:10.1126/scitranslmed.aaf1059

6. Harris SA, Harris EA. Herpes Simplex Virus Type 1 and Other Pathogens are Key Causative Factors in Sporadic Alzheimer’s Disease. Mancuso R, editor. J Alzheimers Dis. 2015;48: 319–53. doi:10.3233/JAD-142853

7. Miklossy J. Bacterial Amyloid and DNA are Important Constituents of Senile Plaques: Further Evidence of the Spirochetal and Biofilm Nature of Senile Plaques. J Alzheimers Dis. 2016;53: 1459–73. doi:10.3233/JAD-160451

8. Corder EH, Saunders AM, Strittmatter WJ, Schmechel DE, Gaskell PC, Small GW, et al. Gene dose of apolipoprotein E type 4 allele and the risk of Alzheimer’s disease in late onset families. Science. 1993;261: 921–3. Available: http://www.ncbi.nlm.nih.gov/pubmed/8346443

9. Cruchaga C, Haller G, Chakraverty S, Mayo K, Vallania FLM, Mitra RD, et al. Rare variants in APP, PSEN1 and PSEN2 increase risk for AD in late-onset Alzheimer’s disease families. PLoS One. 2012;7: e31039. doi:10.1371/journal.pone.0031039

10. Genin E, Hannequin D, Wallon D, Sleegers K, Hiltunen M, Combarros O, et al. APOE and Alzheimer disease: a major gene with semi-dominant inheritance. Mol Psychiatry. 2011;16: 903–7. doi:10.1038/mp.2011.52

11. Guerreiro R, Wojtas A, Bras J, Carrasquillo M, Rogaeva E, Majounie E, et al. TREM2 variants in Alzheimer’s disease. N Engl J Med. 2013;368: 117–27. doi:10.1056/NEJMoa1211851

12. Harold D, Abraham R, Hollingworth P, Sims R, Gerrish A, Hamshere ML, et al. Genome-wide association study identifies variants at CLU and PICALM associated with Alzheimer’s disease. Nat Genet. 2009;41: 1088–93. doi:10.1038/ng.440

13. Hollingworth P, Harold D, Sims R, Gerrish A, Lambert J-C, Carrasquillo MM, et al. Common variants at ABCA7, MS4A6A/MS4A4E, EPHA1, CD33 and CD2AP are associated with Alzheimer’s disease. Nat Genet. 2011;43: 429–35. doi:10.1038/ng.803

14. Jonsson T, Stefansson H, Steinberg S, Jonsdottir I, Jonsson P V, Snaedal J, et al. Variant of TREM2 associated with the risk of Alzheimer’s disease. N Engl J Med. 2013;368: 107–16. doi:10.1056/NEJMoa1211103

15. Lambert JC, Ibrahim-Verbaas CA, Harold D, Naj AC, Sims R, Bellenguez C, et al. Meta-analysis of 74,046 individuals identifies 11 new susceptibility loci for Alzheimer’s disease. Nat Genet. 2013;45: 1452–8. doi:10.1038/ng.2802

16. Naj AC, Jun G, Beecham GW, Wang L-S, Vardarajan BN, Buros J, et al. Common variants at MS4A4/MS4A6E, CD2AP, CD33 and EPHA1 are associated with late-onset Alzheimer’s disease. Nat Genet. 2011;43: 436–41. doi:10.1038/ng.801

17. Mez J, Chung J, Jun G, Kriegel J, Bourlas AP, Sherva R, et al. Two novel loci, COBL and SLC10A2, for Alzheimer’s disease in African Americans. Alzheimers Dement. 2017;13: 119–129. doi:10.1016/j.jalz.2016.09.002

18. Steinberg S, Stefansson H, Jonsson T, Johannsdottir H, Ingason A, Helgason H, et al. Loss-of-function variants in ABCA7 confer risk of Alzheimer’s disease. Nat Genet. 2015;47: 445–7. doi:10.1038/ng.3246

19. Sims R, van der Lee SJ, Naj AC, Bellenguez C, Badarinarayan N, Jakobsdottir J, et al. Rare coding variants in PLCG2, ABI3, and TREM2 implicate microglial-mediated innate immunity in Alzheimer’s disease. Nat Genet. 2017;49: 1373–1384. doi:10.1038/ng.3916

20. Jun GR, Chung J, Mez J, Barber R, Beecham GW, Bennett DA, et al. Transethnic genome-wide scan identifies novel Alzheimer’s disease loci. Alzheimers Dement. 2017;13: 727–738. doi:10.1016/j.jalz.2016.12.012

21. Desikan RS, Fan CC, Wang Y, Schork AJ, Cabral HJ, Cupples LA, et al. Genetic assessment of age-associated Alzheimer disease risk: Development and validation of a polygenic hazard score. Brayne C, editor. PLoS Med. 2017;14: e1002258. doi:10.1371/journal.pmed.1002258

22. Farfel JM, Yu L, Buchman AS, Schneider JA, De Jager PL, Bennett DA. Relation of genomic variants for Alzheimer disease dementia to common neuropathologies. Neurology. 2016;87: 489–496. doi:10.1212/WNL.0000000000002909

23. Auton A, Abecasis GR, Altshuler DM, Durbin RM, Abecasis GR, Bentley DR, et al. A global reference for human genetic variation. Nature. Nature Research; 2015;526: 68–74. doi:10.1038/nature15393

24. Miyashita A, Koike A, Jun G, Wang L-S, Takahashi S, Matsubara E, et al. SORL1 is genetically associated with late-onset Alzheimer’s disease in Japanese, Koreans and Caucasians. Toft M, editor. PLoS One. 2013;8: e58618. doi:10.1371/journal.pone.0058618

25. Karch CM, Ezerskiy LA, Bertelsen S, Goate AM, Goate AM. Alzheimer’s Disease Risk Polymorphisms Regulate Gene Expression in the ZCWPW1 and the CELF1 Loci. Huang Q, editor. PLoS One. 2016;11: e0148717. doi:10.1371/journal.pone.0148717

26. Ryan KJ, White CC, Patel K, Xu J, Olah M, Replogle JM, et al. A human microglia-like cellular model for assessing the effects of neurodegenerative disease gene variants. Sci Transl Med. 2017;9: eaai7635. doi:10.1126/scitranslmed.aai7635

27. GTEx Consortium KG, Deluca DS, Segre A V., Sullivan TJ, Young TR, Gelfand ET, et al. Human genomics. The Genotype-Tissue Expression (GTEx) pilot analysis: multitissue gene regulation in humans. Science. 2015;348: 648–60. doi:10.1126/science.1262110

28. Fritsche LG, Igl W, Bailey JNC, Grassmann F, Sengupta S, Bragg-Gresham JL, et al. A large genome-wide association study of age-related macular degeneration highlights contributions of rare and common variants. Nat Genet. NIH Public Access; 2016;48: 134–43. doi:10.1038/ng.3448

29. Tabata S, Kuroki K, Wang J, Kajikawa M, Shiratori I, Kohda D, et al. Biophysical characterization of O-glycosylated CD99 recognition by paired Ig-like type 2 receptors. J Biol Chem. 2008;283: 8893–901. doi:10.1074/jbc.M709793200

30. Wang J, Fan Q, Satoh T, Arii J, Lanier LL, Spear PG, et al. Binding of herpes simplex virus glycoprotein B (gB) to paired immunoglobulin-like type 2 receptor alpha depends on specific sialylated O-linked glycans on gB. J Virol. 2009;83: 13042–5. doi:10.1128/JVI.00792-09

31. Kogure A, Shiratori I, Wang J, Lanier LL, Arase H. PANP is a novel O-glycosylated PILRa ligand expressed in neural tissues. Biochem Biophys Res Commun. 2011;405: 428–33. doi:10.1016/j.bbrc.2011.01.047

32. Sun Y, Senger K, Baginski TK, Mazloom A, Chinn Y, Pantua H, et al. Evolutionarily conserved paired immunoglobulin-like receptor α (PILRα) domain mediates its interaction with diverse sialylated ligands. J Biol Chem. 2012;287: 15837–50. doi:10.1074/jbc.M111.286633

33. Fournier N, Chalus L, Durand I, Garcia E, Pin JJ, Churakova T, et al. FDF03, a novel inhibitory receptor of the immunoglobulin superfamily, is expressed by human dendritic and myeloid cells. J Immunol. 2000;165: 1197–209. Available: http://www.ncbi.nlm.nih.gov/pubmed/10903717

34. Mousseau DD, Banville D, L’Abbé D, Bouchard P, Shen SH. PILRalpha, a novel immunoreceptor tyrosine-based inhibitory motif-bearing protein, recruits SHP-1 upon tyrosine phosphorylation and is paired with the truncated counterpart PILRbeta. J Biol Chem. 2000;275: 4467–74. Available: http://www.ncbi.nlm.nih.gov/pubmed/10660620

35. Shiratori I, Ogasawara K, Saito T, Lanier LL, Arase H. Activation of Natural Killer Cells and Dendritic Cells upon Recognition of a Novel CD99-like Ligand by Paired Immunoglobulin-like Type 2 Receptor. J Exp Med. 2004;199: 525–533. doi:10.1084/jem.20031885

36. Wang J, Shiratori I, Uehori J, Ikawa M, Arase H. Neutrophil infiltration during inflammation is regulated by PILRα via modulation of integrin activation. Nat Immunol. 2013;14: 34–40. doi:10.1038/ni.2456

37. Sun Y, Caplazi P, Zhang J, Mazloom A, Kummerfeld S, Quinones G, et al. PILRα Negatively Regulates Mouse Inflammatory Arthritis. J Immunol. 2014;193: 860–870. doi:10.4049/jimmunol.1400045

38. Kohyama M, Matsuoka S, Shida K, Sugihara F, Aoshi T, Kishida K, et al. Monocyte infiltration into obese and fibrilized tissues is regulated by PILRα. Eur J Immunol. 2016;46: 1214–1223. doi:10.1002/eji.201545897

39. Kuroki K, Wang J, Ose T, Yamaguchi M, Tabata S, Maita N, et al. Structural basis for simultaneous recognition of an O-glycan and its attached peptide of mucin family by immune receptor PILRα. Proc Natl Acad Sci U S A. 2014;111: 8877–82. doi:10.1073/pnas.1324105111

40. Halim A, Rüetschi U, Larson G, Nilsson J. LC-MS/MS characterization of O-glycosylation sites and glycan structures of human cerebrospinal fluid glycoproteins. J Proteome Res. 2013;12: 573–84. doi:10.1021/pr300963h

41. Lu Q, Lu G, Qi J, Wang H, Xuan Y, Wang Q, et al. PILRα and PILRβ have a siglec fold and provide the basis of binding to sialic acid. Proc Natl Acad Sci U S A. 2014;111: 8221–6. doi:10.1073/pnas.1320716111

42. Satoh T, Arii J, Suenaga T, Wang J, Kogure A, Uehori J, et al. PILRalpha is a herpes simplex virus-1 entry coreceptor that associates with glycoprotein B. Cell. 2008;132: 935–44. doi:10.1016/j.cell.2008.01.043

43. Srinivasan K, Friedman BA, Larson JL, Lauffer BE, Goldstein LD, Appling LL, et al. Untangling the brain’s neuroinflammatory and neurodegenerative transcriptional responses. Nat Commun. 2016;7: 11295. doi:10.1038/ncomms11295

44. Wang Y, Cella M, Mallinson K, Ulrich JD, Young KL, Robinette ML, et al. TREM2 Lipid Sensing Sustains the Microglial Response in an Alzheimer’s Disease Model. Cell. 2015;160: 1061–1071. doi:10.1016/j.cell.2015.01.049

45. Orre M, Kamphuis W, Osborn LM, Jansen AHP, Kooijman L, Bossers K, et al. Isolation of glia from Alzheimer’s mice reveals inflammation and dysfunction. Neurobiol Aging. 2014;35: 2746–2760. doi:10.1016/j.neurobiolaging.2014.06.004

46. Wyss-Coray T, Yan F, Lin AH-T, Lambris JD, Alexander JJ, Quigg RJ, et al. Prominent neurodegeneration and increased plaque formation in complement-inhibited Alzheimer’s mice. Proc Natl Acad Sci U S A. 2002;99: 10837–42. doi:10.1073/pnas.162350199

47. Lambert J-C, Heath S, Even G, Campion D, Sleegers K, Hiltunen M, et al. Genome-wide association study identifies variants at CLU and CR1 associated with Alzheimer’s disease. Nat Genet. 2009;41: 1094–1099. doi:10.1038/ng.439

48. Bouchon A, Hernández-Munain C, Cella M, Colonna M. A DAP12-mediated pathway regulates expression of CC chemokine receptor 7 and maturation of human dendritic cells. J Exp Med. 2001;194: 1111–22. Available: http://www.ncbi.nlm.nih.gov/pubmed/11602640

49. Colonna M, Wang Y. TREM2 variants: new keys to decipher Alzheimer disease pathogenesis. Nat Rev Neurosci. 2016;17: 201–7. doi:10.1038/nrn.2016.7

50. Alam M, Alam Q, Kamal M, Jiman-Fatani A, Azhar E, Khan M, et al. Infectious Agents and Neurodegenerative Diseases: Exploring the Links. Curr Top Med Chem. 2017;17: 1390–1399. doi:10.2174/1568026617666170103164040

51. Itzhaki RF, Lin WR, Shang D, Wilcock GK, Faragher B, Jamieson GA. Herpes simplex virus type 1 in brain and risk of Alzheimer’s disease. Lancet (London, England). 1997;349: 241–4. doi:10.1016/S0140-6736(96)10149-5

52. Steel AJ, Eslick GD. Herpes Viruses Increase the Risk of Alzheimer’s Disease: A Meta-Analysis. J Alzheimer’s Dis. 2015;47: 351–364. doi:10.3233/JAD-140822

53. Lövheim H, Gilthorpe J, Johansson A, Eriksson S, Hallmans G, Elgh F. Herpes simplex infection and the risk of Alzheimer’s disease: A nested case-control study. Alzheimers Dement. 2015;11: 587–92. doi:10.1016/j.jalz.2014.07.157

54. Letenneur L, Pérès K, Fleury H, Garrigue I, Barberger-Gateau P, Helmer C, et al. Seropositivity to herpes simplex virus antibodies and risk of Alzheimer’s disease: a population-based cohort study. Hornung R, editor. PLoS One. 2008;3: e3637. doi:10.1371/journal.pone.0003637

55. Mancuso R, Baglio F, Cabinio M, Calabrese E, Hernis A, Nemni R, et al. Titers of herpes simplex virus type 1 antibodies positively correlate with grey matter volumes in Alzheimer’s disease. J Alzheimers Dis. 2014;38: 741–5. doi:10.3233/JAD-130977

56. Agostini S, Mancuso R, Baglio F, Cabinio M, Hernis A, Costa AS, et al. High avidity HSV-1 antibodies correlate with absence of amnestic Mild Cognitive Impairment conversion to Alzheimer’s disease. Brain Behav Immun. 2016;58: 254–260. doi:10.1016/j.bbi.2016.07.153

57. Huemer HP, Menzel HJ, Potratz D, Brake B, Falke D, Utermann G, et al. Herpes simplex virus binds to human serum lipoprotein. Intervirology. 1988;29: 68–76. Available: http://www.ncbi.nlm.nih.gov/pubmed/2842273

58. Wozniak MA, Itzhaki RF, Shipley SJ, Dobson CB. Herpes simplex virus infection causes cellular beta-amyloid accumulation and secretase upregulation. Neurosci Lett. 2007;429: 95–100. doi:10.1016/j.neulet.2007.09.077

59. De Chiara G, Marcocci ME, Civitelli L, Argnani R, Piacentini R, Ripoli C, et al. APP processing induced by herpes simplex virus type 1 (HSV-1) yields several APP fragments in human and rat neuronal cells. Blaho JA, editor. PLoS One. 2010;5: e13989. doi:10.1371/journal.pone.0013989

60. Piacentini R, Civitelli L, Ripoli C, Marcocci ME, De Chiara G, Garaci E, et al. HSV-1 promotes Ca2+-mediated APP phosphorylation and Aβ accumulation in rat cortical neurons. Neurobiol Aging. 2011;32: 2323.e13–26. doi:10.1016/j.neurobiolaging.2010.06.009

61. Sokolov AA, Reincke M. Herpes simplex encephalitis affecting the entire limbic system. Mayo Clin Proc. 2012;87: e69. doi:10.1016/j.mayocp.2012.06.023

62. Lathe R, Haas JG. Distribution of cellular HSV-1 receptor expression in human brain. J Neurovirol. Springer; 2017;23: 376–384. doi:10.1007/s13365-016-0504-x

63. Carter CJ. APP, APOE, complement receptor 1, clusterin and PICALM and their involvement in the herpes simplex life cycle. Neurosci Lett. 2010;483: 96–100. doi:10.1016/j.neulet.2010.07.066

64. Huemer HP, Wang Y, Garred P, Koistinen V, Oppermann S. Herpes simplex virus glycoprotein C: molecular mimicry of complement regulatory proteins by a viral protein. Immunology. 1993;79: 639–47. Available: http://www.ncbi.nlm.nih.gov/pubmed/8406590

65. Blaho JA, Morton ER, Yedowitz JC. Herpes simplex virus: propagation, quantification, and storage. Current protocols in microbiology. Hoboken, NJ, USA: John Wiley & Sons, Inc.; 2005. p. Unit 14E. 1. doi:10.1002/9780471729259.mc14e01s00

